# Functional and topological analysis of PSENEN, the fourth subunit of the γ-secretase complex

**DOI:** 10.1101/2023.07.28.550932

**Authors:** Lutgarde Serneels, Leen Bammens, An Zwijsen, Alexandra Tolia, Lucía Chávez-Gutiérrez, Bart De Strooper

**Author notes:** Address correspondence to: Bart De Strooper, Herestraat 49 3000 Leuven, Belgium.

## Abstract

The γ-secretase complexes are intramembrane cleaving proteases involved in the generation of the Aβ peptides in Alzheimer’s Disease. The complex consists of four subunits, with Presenilin, harboring the catalytic site. Here, we study the role of the smallest subunit, PSENEN or Presenilin enhancer 2 (PEN-2), encoded by the gene *Psenen*, *in vivo* and *in vitro*. We find a profound Notch-deficiency phenotype in *Psenen^−/−^* embryos confirming the essential role of PSENEN in the γ-secretase complex. We used *Psenen^−/−^* fibroblasts to explore the structure-function of PSENEN by the Scanning Cysteine Accessibility Method. Glycine22 and Proline27 which border the membrane domains 1 and 2 of PSENEN are involved in complex formation and stabilization of γ-secretase. The hairpin structured hydrophobic membrane domains 1 and 2 are exposed to a water-containing cavity in the complex, while transmembrane domain 3 is not water exposed. We finally demonstrate the essential role of PSENEN for the cleavage activity of the complex. PSENEN is more than a structural component of the γ-secretase complex and might contribute to the catalytic mechanism of the enzyme.

## Introduction

γ-Secretase is a membrane-embedded aspartic protease composed of four subunits: Presenilin 1 or 2 (PSEN1 or PSEN2), Nicastrin (NCSTN), APH1A or APH1B, and PSENEN (1). γ-Secretase is an attractive drug target for Alzheimer’s disease (AD) because it is responsible for the final cleavage in the processing of the Amyloid Precursor Protein (APP) to Aβ peptides(2). However, due to its broad substrate specificity, general inhibition of γ-secretase is not without risks. The main problem is the involvement of γ-secretase in the cleavage of Notch (3–5), which releases the Notch Intracellular Domain (NICD) that is implicated in crucial signaling processes throughout life. The essential role of γ-secretase in early development is illustrated by the embryonic lethality of *D. melanogaster*, *C. elegans* or *M. musculus* in which *Ncstn*, *Aph1A,* or the two *Psens* together are genetically inactivated. In all cases a severe Notch-deficiency phenotype is observed (5–15). However, it should be noted that other single knockouts of subunits of the γ-secretase complex, like *Psen2* alone (7, 10) or *Aph1B* (8, 16) do not result in Notch phenotypes. It is also increasingly clear that γ-secretases play a role in various other signaling pathways (17). The effect of a genetic knockout of *PSENEN* causing a Notch deficient phenotype is reported here, confirming our retracted report (18).

The elucidation of the structure of γ-secretase (19) has provided a quantum leap in our understanding of these enzymes. The tetrameric γ-secretase encompasses twenty transmembrane domain helices, organised in a horseshoe conformation (20). The catalytic aspartates D257 and D385 are located on transmembrane domain (TM) 6 and TM7 (21) of the PSEN subunit. The large ectodomain of NCSTN overlays the whole hydrophobic domain of γ-secretase at the extracellular surface. The catalytic aspartates are present in a water containing cavity to which other parts of PSEN contribute as well (22–24). The role of the other three subunits of γ-secretase and their contribution to the catalytic activity of the protease are less well understood, although NCSTN is involved in substrate binding (25). All subunits are needed to generate a mature tetrameric γ-secretase complex. In the current work we focus on the smallest subunit, presenilin enhancer 2 or PSENEN, previously called PEN-2.

*Psenen* was originally identified in a genetic screen for modulators of PSEN activity in *C. elegans* (11). PSENEN is a 101 amino acids long protein with three hydrophobic domains. Initially several publications suggested a hairpin topology for the protein, with the loop domain exposed to the intracellular side of the cell membrane (18, 26, 27). The cryo-EM structure of γ-secretase (20) has shown however that the two first hydrophobic domains of PSENEN make a re-entry hairpin loop, resulting in the exposure of its N-terminus to the cytoplasmic side of the membrane and the C-terminus to the luminal or extracellular side of the membrane. This was also confirmed by other approaches (28).

RNAi mediated down regulation of PSENEN levels in cell culture leads to decreased endo-proteolysis of PSEN, which is associated with an increase of full length (FL) PSEN and a decrease of PSEN amino- and carboxy-terminal fragments (PSEN NTF and PSEN CTF) (29–32). Additionally, mutational analysis has shown that the N-terminal part of hydrophobic domain 1 of PSENEN interacts with the TMD4 of PSEN1 and is important for PSEN endo-proteolysis (31, 33). These observations suggest that PSENEN is involved in the endo-proteolysis of PSEN and therefore in the activation of the γ-secretase complex (34). Furthermore, a conserved amino acid sequence motif, DYSLF, in the carboxy-terminus of PSENEN, as well as the length of this part of the protein, appear crucial for the assembly of the γ-secretase complex, the stabilization of the PSEN fragments after endo-proteolysis and for γ-secretase activity (31, 35–37). Incorporation of a Flag-tag at the N-terminus of PSENEN changes the conformation of PSEN, resulting in an increased Aβ_42_/Aβ_40_ ratio (38), similar to what is observed for familial Alzheimer’s disease mutations in the PSEN subunit (15). Furthermore, a γ-secretase modulator that decreases Aβ_42_ production binds mainly to PSENEN (39), further arguing for the crucial role of PSENEN in the regulation of the activity of the complex.

We confirm the conclusions from our previous report (18) that PSENEN is essential for the γ-secretase processing of Notch and APP. In the absence of PSENEN, no processing of APP is observed, which contrasts with *Ncstn^−/−^* cells, which retain 5-6% of γ-secretase activity (40). We made use of the *Psenen^−/−^* fibroblasts to perform a cysteine scanning analysis (41) of the PSENEN protein (22–24) to investigate its topology and the exposure of individual amino acids to water.

## EXPERIMENTAL PROCEDURES

### Generation of Psenen^−/−^ embryos

*Psenen^+/−^* male and female mice were purchased from the Texas Institute for Genomic Medicine and coupled to obtain *Psenen^−/−^* embryos. Mice were kept on a C57Bl6J background and housed in cages enriched with woodwool and shavings as bedding and given access to water and food ad libitum. All experiments were approved by the Ethical Committee for Animal Experimentation at the University of Leuven (KU Leuven).

### Whole mount in situ hybridization

Embryos were isolated from the mother at the indicated embryonic days and fixed in 4% paraformaldehyde in PBS. After dehydration to 100% methanol and bleaching in 6% hydrogen peroxide for 1 hour at room temperature, embryos were rehydrated in 100% PBS + 0.1% Tween. The embryos were treated with 10 µg/ml proteinase K for 5 min and post-fixated in 0.2% gluteraldehyde and 4% paraformaldehyde. Prehybridization was done in 50% formamide and 5 SSC pH 4.5 complemented with 50 µg/ml yeast RNA and 50 µg/ml porcine heparine during 1 hour at 60°C, after which the prehybridisation mix was replaced for the hybridization mix containing 1ng/µl digitonin-labeled riboprobes against Notch signaling pathway components as indicated. After overnight incubation in the hybridization mix at 60°C, the embryos were washed extensively in PBS with 0.1% Tween and treated with RNase A (100mg/ml) for 15 min.

Blocking was performed in 2% Boehringer Blocking Reagent and 20% fetal bovine serum in MABT buffer (100mM Maleic Acid, 150mM NaCl, pH 7.5, 0.1% Tween). After blocking, the embryos were treated with an alkaline phosphatase-coupled anti-digitonin antibody (final concentration 1/2000) overnight at 4°C. The next day, the embryos were washed in MABT and treated with levamisole (2mM). BM purple, a chromogenic substrate for alkaline phosphatase, was added and the staining was visualized by light microscopy.

### Generation of Psenen^−/−^ fibroblasts and cell culture

Fibroblasts with the different genotypes as indicated, were generated from E9.5 embryos and immortalized by transfection with the plasmid pMSSVLT, driving expression of the large T-antigen. Immortalized mouse embryonic fibroblasts (MEFs) were cultured in Dulbecco’s modified Eagle’s medium/F-12 containing 10% fetal bovine serum (Sigma).

### Generation of PSENEN mutants and corresponding stable cell lines

All mutations in PSENEN were generated with the XL site-directed-mutagenesis kit (Stratagene) and confirmed by DNA sequence analysis. Retroviruses were generated by co-transfecting pMSCVpuro vector containing PSENEN and the PIK helper plasmid into HEK293 cells. Viral particles harvested at 48h post-transfection were used to infect *Psenen^−/−^* fibroblasts at 30-40% confluency. Transduced cells were selected with 5 µg/ml puromycin.

### Antibodies

Polyclonal antibodies against mouse PSENEN (B126.2), PSEN1 NTF (B19.3), APH1A (B80.2) and APP C-terminus (B63.3), and monoclonal 9C3 against the C-terminus of NCTSN have been described previously (42, 43). The following antibodies were purchased: anti-N-cadherin from BD Biosciences, anti-NICD (cleaved Notch1 Val-1744) from Cell Signaling Technologies, mAb 9E10 (Sanver Tech), and MAB5232 against PSEN1 CTF (Chemicon). The anti-Flag M2 antibody was purchased from Sigma-Aldrich.

### SDS-PAGE

Total cell lysates were prepared in lysis buffer (5 mM Tris-HCl pH 7.4, 250 mM sucrose, 1 mM EGTA, 1% Triton X-100, and complete protease inhibitors (Roche Applied Science)). Post-nuclear fractions were taken and protein concentrations were determined using standard BCA assay (Pierce). 20 µg of protein was separated on 4-12% BisTris gels and transferred to nitrocellulose membranes to perform western blot analysis. Signals were detected using the chemiluminiscence detection with Renaissance (PerkinElmer) and developed either by X-ray film or LAS-3000 (Fuji). Quantifications of the western blots was done with Fiji (Image J).

### Blue Native PAGE

Microsomal membrane fractions were prepared in lysis buffer containing 0.5% dodecylmaltoside, 20% glycerol and 25% BisTris/HCl pH7. Upon ultracentrifugation (55000 rpm), supernatant was taken, protein concentrations were measured, and 5 µg of protein was supplemented with sample buffer. Samples were loaded on a 5-16% polyacrylamide gel and run for 4 hours at 4°C. Subsequently, the gel was incubated with 0.1 % SDS for 10 min at room temperature and transferred to a polyvinylidene difluoride membrane. Membranes were de-stained in water/methanol/acetic acid (60/30/10, v/v/v) and incubated with antibodies to detect γ-secretase complex.

### Water accessibility assay

Cells were plated in 10 cm dishes, washed with PBS, and treated with the biotinylated sulfhydryl-specific reagents for 30 min at 4°C. In case of pretreatment with Sodium (2-Sulfonatoethyl) methanethiosulfonate (MTSES), cells were incubated with MTSES or DMSO for 30 min at 4°C before treatment with the biotinylated reagent. After extensive washing in PBS to remove unbound reagent, cells were collected and lysed in 25 mM HEPES pH8, 150 mM NaCl, 2 mM EDTA, 1% Triton X-100. Biotinylated proteins in the total cell lysates were precipitated with immobilized NeutrAvidin protein beads (Pierce). Bound proteins were eluted by boiling in Nu-Page sample buffer and SDS-PAGE was performed as mentioned above. PSENEN was detected in input and bound fractions.

Sulfhydryl-specific reagents used: EZ-Link Biotin-HPDP (N-(6-(Biotinamido)hexyl)-3’-(2’-pyridyldithio)-propionamide, Pierce) (200 µM), MTSEA-biotin (N-biotinylaminoethyl-methanethiosulfonate, Biotium) (500 µM) and TS-XX-biotin ethylenediamine (biotin-XX ethylenediamine thiosulfate, sodium salt, Biotium) (500 µM) and MTSES (Sodium (2-Sulfonatoethyl)methanethiosulfonate, Biotium) (200 µM).

### Crosslinking

Crosslinking studies were performed by incubating microsomal membrane fractions with the heterobifunctional amine-sulfhydryl-reactive cross linker SPDP (*N*-Succinimidyl-3-(2-pyridyldithio)-propionate) (1 mM). As negative controls, membranes preincubated with 10 mM N-ethylmaleimide (NEM)/10 mM EDTA or membranes incubated with DMSO were used.

Reactions were quenched with NEM/EDTA. After centrifugation at 55000 rpm, membrane pellets were solubilized in 5 mM Tris-HCl pH 7.4, 250 mM sucrose, 1 mM EGTA, 1% Triton X-100, equal amounts of proteins were separated in SDS-PAGE in non-reducing conditions and detected by western blotting using antibodies against γ-secretase components.

### Cell free APP processing assay

Cell free assays were performed as described by Kakuda *et al*. with some minor modifications (44). Briefly, microsomal membrane fractions solubilized in 1% CHAPSO were mixed with recombinant APPC99-3xFlag substrate (0.5 µM final concentration), 0.0125% phosphatidylethanolamine, 0.1% phosphatidylcholine and 2.5% DMSO. Reactions were incubated at 37°C for 3 hours. AICD was detected by western blot analysis with the anti-Flag M2 antibody and Aβ species by the AlphaLISA technique (see below).

Aβ and AICD levels were normalized to the amounts of γ-secretase complex in the *in vitro* assay, which were estimated from the PSEN1 NTF levels.

### Cell based APP processing assay

Fibroblasts were infected with an adenoviral vector (Ad5) bearing human APP-695 containing the Swedish mutation. The cells were then cultured in Dulbecco’s modified Eagle’s medium supplemented with 0.2% fetal bovine serum for 16 hours, and the conditioned medium was collected and used to analyze APP processing. Aβ_40_ and Aβ_42_ levels were quantified by AlphaLISA (see below) and sAPP levels by SDS-PAGE and western blot analysis. Cell lysates were prepared and APP full length and APP CTF fragments were analyzed by SDS-PAGE followed by western blot analysis. Aβ levels were normalized to infection efficiency, quantified from the sAPP expression levels. APP CTF were normalized to APP full length.

### Cell based Notch processing assay

Fibroblasts were infected with an adenoviral vector (Ad5) containing Myc-tagged NotchΔE. At 24 hours post-infection, 10 µM lactacystin was added to the cultures and after 4 hours cell lysates were prepared. NICD and NotchΔE levels were estimated by Western Blot analysis using a neo-epitope (cleaved Notch1 Val-1744) and an anti-myc antibody, respectively. NICD levels were normalized to the levels of infection efficiency (levels of NotchΔE).

### AlphaLISA (Perkin Elmer)

Conditioned medium was mixed with PBS supplemented with 0.1% caseine, streptavidine-coated AlphaLISA donor beads and biotinylated antibody against the neo-epitope of Aβ_40_ or Aβ_42_. After overnight incubation at 4°C, acceptor-beads coupled to an antibody against the N-terminus of Aβ were added. After 1 hour incubation, light emission (615 nm) was detected upon laser excitation at 680 nm.

### Statistical analysis

Data from 4 experiments were used for calculation of p-values using one-way ANOVA and Turkey multiple comparisons test.

## RESULTS AND DISCUSSION

### *Psenen^−/−^* embryos display a Notch-deficiency phenotype

We generated a *Psenen^−/−^* mouse model by crossing *Psenen^+/−^* mice obtained from the Texas Institute for Genomic Medicine. The *Psenen* gene was inactivated by exchanging three of the four *Psenen* exons (exon 2, 3, and 4) for a *LacZ/Neomycin* cassette. *Psenen* gene disruption resulted in embryonic lethality, confirming the essential role of PSENEN in the maturation and/or stabilization of the γ-secretase complex. Up to E9.5, living *Psenen*^−/−^ embryos were recovered in a nearly Mendelian ratio (21/93), while they were completely absorbed by E11.

At E9.5, the yolk sac surrounding the *Psenen^−/−^* embryos had only a primitive immature blood vessel-plexus and a blistered appearance compared to the yolk sac of the *Psenen^+/+^* or *Psenen^+/−^* littermates, which displayed a normal complex vasculature (Fig. 1A). The *Psenen*^−/−^ embryos were smaller than the *Psenen^+/+^* littermates. They had an abnormally large pericardial sac, a kinked neural tube, and a truncated posterior region. Somitogenesis was initiated in the *Psenen^−/−^* embryos but appeared severely delayed compared to the *Psenen^+/+^* littermates. The optic and otic vesicles, the first branchial arch and the forelimb buds were visible, but the fusion of the headfolds was delayed (Fig. 1A).

**Fig. 1:**
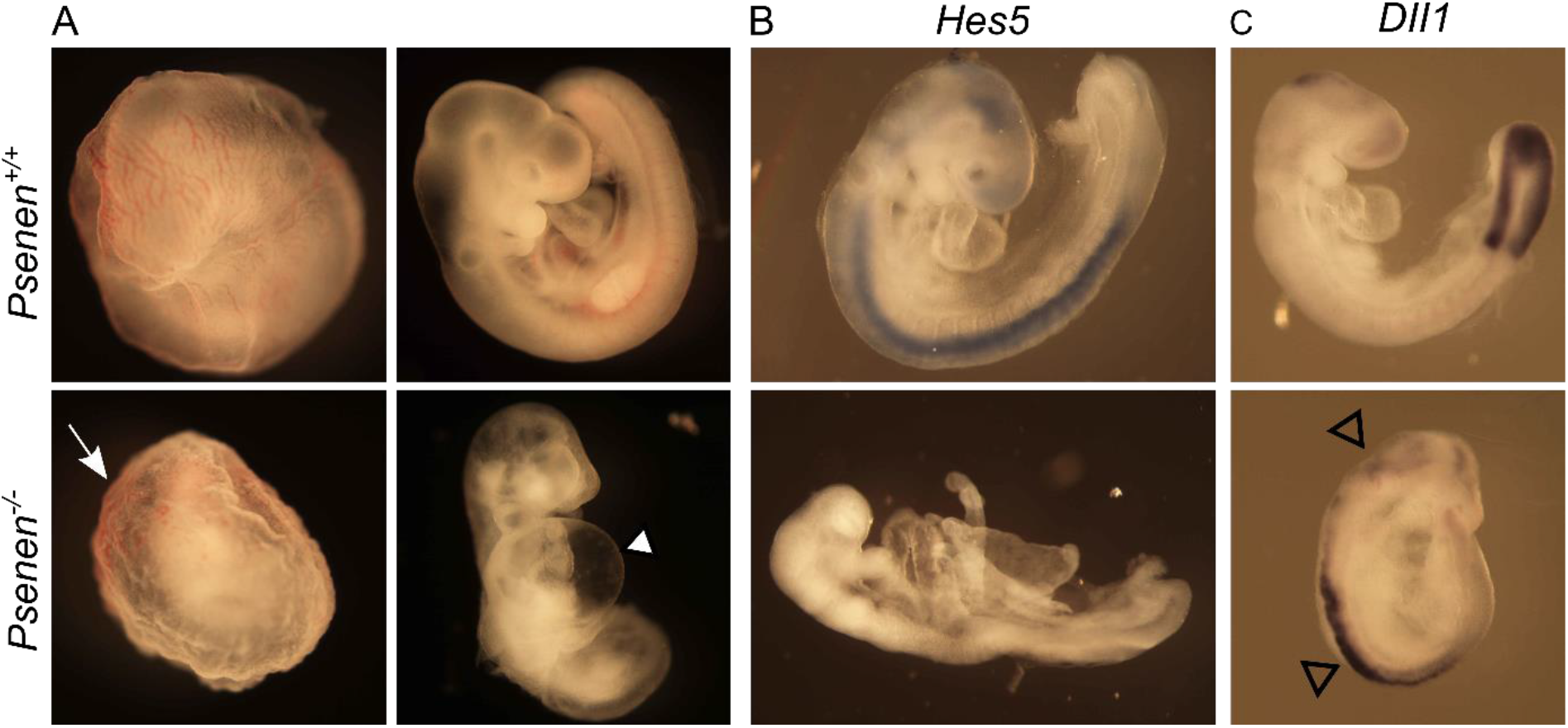
Notch-deficiency phenotype in *Psenen^−/−^* embryos. **(A)** The yolk sac (E9.5) of a *Psenen^+/+^* has a smooth appearance with a hierarchically organized vascular network (upper left panel). In *Psenen^−/−^* (lower panel), an initial vascular plexus and primitive red blood cells have formed (arrow) in the yolk sac, but organization into a discrete network of vitelline vessels is lacking. *Psenen^−/−^* embryos (E9) are developmentally delayed. The knockout embryos are smaller than wildtype littermates and characterized by a posterior truncation, a large pericardial sac (solid arrowhead) and a distorted neural tube. **(B, C)** mRNA expression of two target genes of Notch, *Hes5* and *Dll1,* detected by whole mount *in situ* hybridization with digitonin-labeled riboprobes (blue-brownish color) shows disturbed Notch signaling in *Psenen^−/−^* embryos. The *Hes5* probe (B) reveals repressed *Hes5* expression in *Psenen^−/−^* embryos (E8.5) compared to *Psenen^+/+^* embryos (E8.5) going from the head to the tail along the spinal cord. *Dll1* (C) is ectopically expressed in the neural tube and the head in *Psenen^−/−^* embryos (open arrowheads).

The phenotype of the *Psenen^−/−^* embryos was similar to the previously characterized phenotype of the double knock-out (dKO) *Psen1&2^dKO^* embryos and is compatible with loss of function of γ-secretase activity and the consequent effects on the Notch signaling pathways (7, 10). To evaluate disturbance in Notch signaling, we performed whole mount *in situ* hybridization on embryos at E8.5 using *Hes-5* and *Delta-like1* probes to detect expression of the respective genes. If Notch cleavage by γ-secretase is blocked, then *Hes-5* signals are expected to be down- and *Delta-like1* signals to be up-regulated, respectively. We observed indeed the absence of *Hes-5* mRNA and ectopic expression of *Delta-like-1* mRNA in *Psenen^−/−^* compared to the *Psenen^+/+^* littermate embryos (Fig. 1B and C).

The data confirm that PSENEN is an essential component of γ-secretase. Due to the severe Notch loss-of-function phenotype, it is difficult to detect potential non-Notch signaling dependent phenotypes. This is like animals with the other γ-secretase components inactivated (7–10, 16). This precluded assessing the contribution of other γ-secretase substrates in the *Psenen^−/−^* phenotype and to rule out the possibility that PSENEN has additional non-γ-secretase related functions. However, considering that PSENEN protein expression is strongly dependent on the presence of the three other γ-secretase components (8, 16, 32, 45), it seems logical to assume that PSENEN mainly functions as part of the γ-secretase complex.

### Effect of PSENEN deficiency on γ-secretase activity and assembly

We derived *Psenen^+/+^*, *Psenen*^+/−^ and *Psenen^−/−^* fibroblasts from mouse embryos at E9.5 and immortalized the cells. Western blot analysis of *Psenen^−/−^* cell membrane extracts demonstrated the complete absence of PSENEN protein. In contrast, the three other components of the γ-secretase complex were present at normal levels (i.e., equal to the expression in *Psenen^+/+^* cells) (Fig. 2A). Interestingly, PSEN1 was mainly present as full-length protein and NCSTN failed to become fully glycosylated. We observed an accumulation of APP carboxyterminal fragments (CTF), indicating the γ-secretase activity deficiency in the *Psenen^−/−^* fibroblasts (Fig. 2A). γ-Secretase maturation and activity correlated with the expression levels of PSENEN in these cells as assessed in *Psenen^+/−^* cells, suggesting a limiting role of PSENEN in the assembly and activity of the complex in this cell type (Fig. 2A and 2B).

**Fig. 2:**
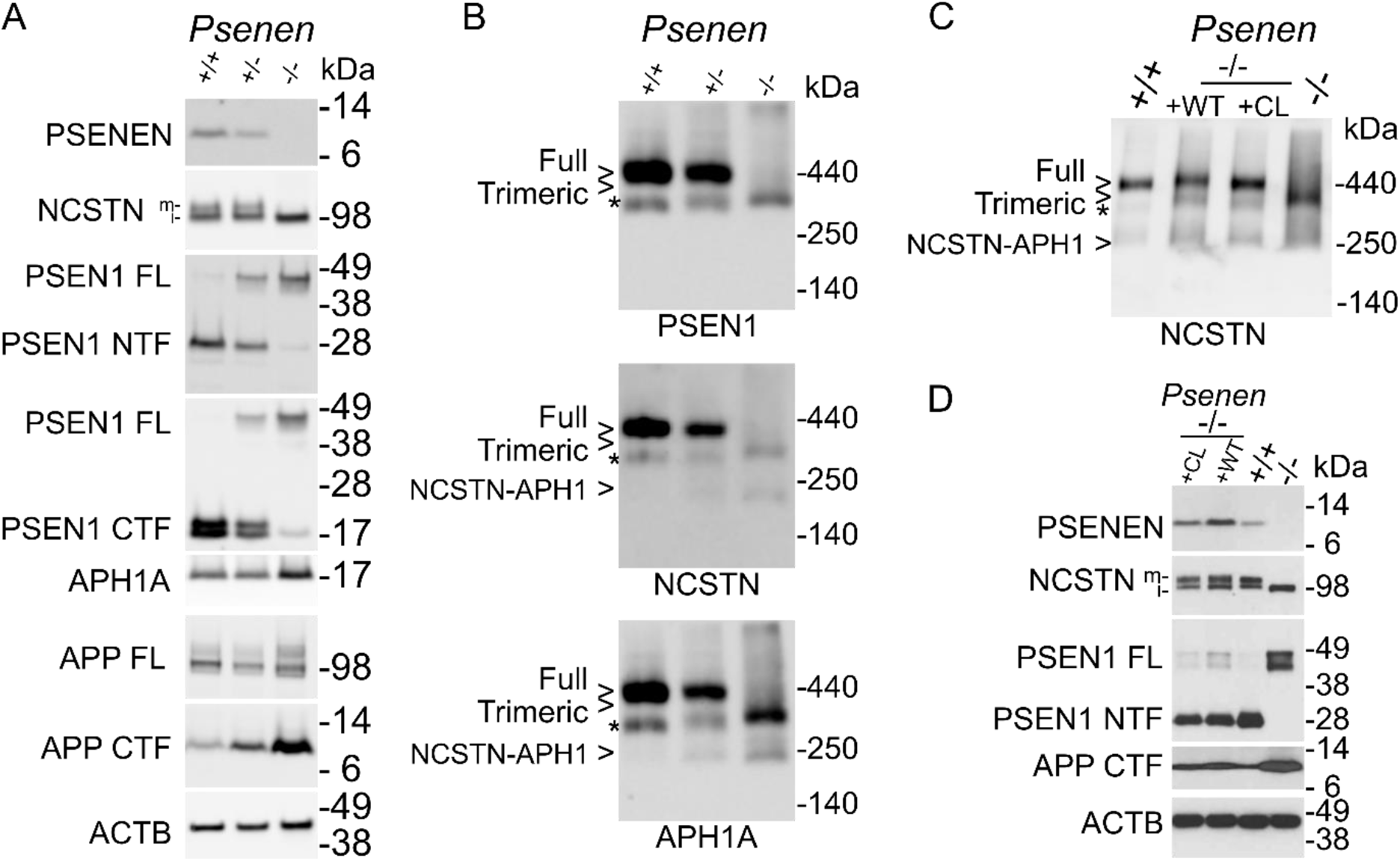
Characterization of *Psenen^+/−^* and *Psenen^−/−^* fibroblasts. **(A)** 4-12% BisTris SDS-PAGE/western blot analysis of *Psenen^+/+^*, *Psenen^+/−^* and *Psenen^−/−^* fibroblasts derived from E9.5 mice. 20 µg solubilized membrane protein was applied per lane and two gels were run and transferred to nitrocellulose membranes. Based on molecular weight markers the membranes were cut and stained with the indicated antibodies. Membrane 1 was stained with antibody B128 to show PSENEN, with antibody 9C3 to demonstrate immature (i) and mature (m) NCSTN, and Mab5232 to reveal full length Presenilin 1 (PSEN FL) and C-terminal fragments (PSEN1 CTF). Membrane 2 was stained with B19 to reveal PSEN1 FL and Presenilin1 N-terminal fragments (PSEN1 NTF), B63 was used to stain full length APP (APP FL) and APP C-terminal fragments (APP CTF) and B80 for APH1A. ACTB was used as loading control (result of membrane 2 is presented here). **(B)** Blue Native PAGE/western blot analysis of *Psenen^+/+^*, *Psenen^+/−^* and *Psenen^−/−^* DDM extracted membranes (5 µg/lane) stained with antibodies against PSEN1 CTF, NCSTN and APH1A as in panel A. Full γ-secretase complex is absent in *Psenen^−/−^* fibroblasts. Two gels were loaded, membrane 1 was first stained with Mab5232 against PSEN1 CTF (upper panel) and reprobed with B80 against APH1A (lower panel) while membrane 2 was stained with 9C3 against NCSTN (middle panel). The trimeric complex in *Psenen^−/−^* fibroblasts is composed of FL PSEN1, NCSTN and APH1A. * indicates a complex induced by detergent-dependent dissociation, composed of PSEN1 CTF, APH1A and NCSTN, see ref (66) which moves slightly faster than the trimeric complex in *Psenen^−/−^*. **(C)** Blue Native PAGE/western blot analysis (5 µg/lane) of membrane fractions of *Psenen^+/+^* fibroblasts, of *Psenen^−/−^* fibroblasts reconstituted with wild type PSENEN (WT) or cysteine-less PSENEN (CL) and *Psenen^−/−^* fibroblasts. The cysteine-less PSENEN mutant is incorporated in the mature γ-secretase complex to the same extent as the wild type PSENEN. *: complex induced by detergent (see panel A). **(D)** SDS-PAGE/western blot analysis of membranes of fibroblasts as in panel C and using the same antibodies as in panel A. The cysteine-less PSENEN mutant rescues NCSTN maturation, PSEN1 endo-proteolysis and restores the cleavage of APP CTF to the same extent as wild type PSENEN. Samples were loaded on two gels. The first membrane was used to detect PSENEN and ACTB. The second membrane was stained for NCSTN, PSEN1 FL (full length) and NTF (amino-terminal fragment). Afterwards this blot was probed for APP CTF.

In Blue Native PAGE analysis, a trimeric complex was detected in *Psenen^−/−^* fibroblasts which is composed of full length PSEN1, NCSTN, and APH1A (Fig. 2B). Since this trimeric complex contains the catalytic subunit of γ-secretase (PSEN), we wondered whether this subcomplex had some remaining enzymatic activity. We measured Aβ levels in the conditioned medium of *Psenen^+/+^* and *Psenen^−/−^* fibroblasts using the AlphaLISA technology (Fig. 3C and E) showing that Aβ_40_ and, Aβ_42_ generation was abolished. We also tested the activity of the complex in a cell free assay by solubilizing microsomal membrane fractions of the *Psenen^+/+^* and *Psenen^−/−^* fibroblasts in 1% CHAPSO and adding recombinant APP-3XFlag substrate (Fig. 3D). AICD levels in the *in vitro* reactions were analysed by SDS-PAGE and western blotting. This experiment showed that AICD production was reduced to background levels (Fig 3D). Western blot analysis showed that in the *Psenen^−/−^* fibroblasts APP CTF fragments accumulated (Fig. 2A,D and 3C). Finally, a dilution series of conditioned medium of *Psenen^+/+^* fibroblasts demonstrated that Aβ_40_ production in *Psenen^−/−^* fibroblasts was less than 1% of the Aβ_40_ production in *Psenen^+/+^* fibroblasts (Fig.3E). This contrasts with the higher activity in Aβ production from *Ncstn^−/−^* fibroblasts, which was > 3% of the wild-type production. This result confirms previous work that showed that in the absence of NCSTN some γ-secretase activity can still be measured (40). PSENEN on the other hand, appears to be necessary to generate an active complex in cell culture. This is in line with an in vitro reconstitution experiment that demonstrated that PSENEN is needed to activate (wild type) PSEN(34). AICD production was also completely abolished indicating that not only the γ-secretase cleavage of APP was disturbed, but the ε-cleavage as well (Fig 3D).

**Fig. 3:**
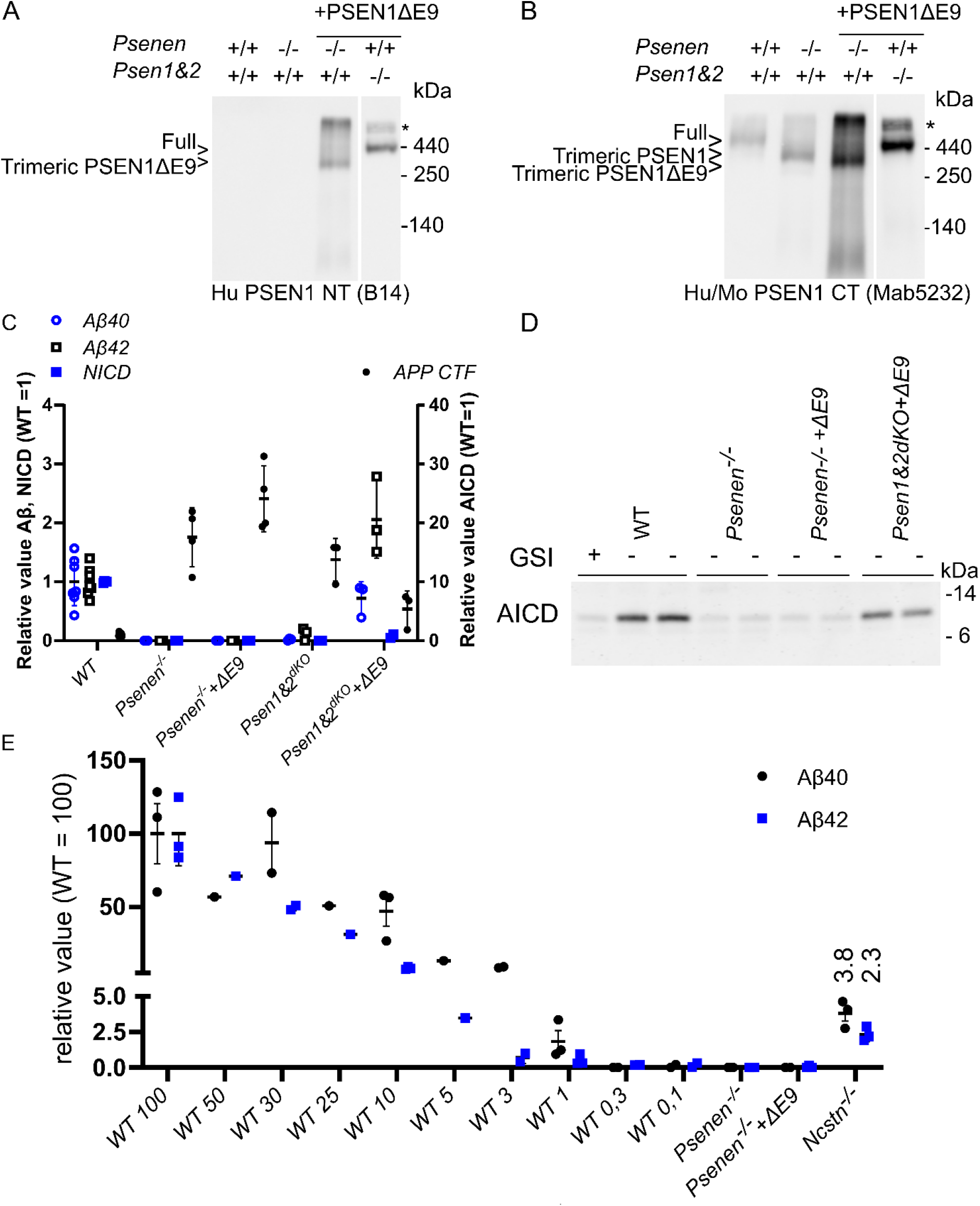
The lack of γ-secretase activity in *Psenen^−/−^* fibroblasts is not simply caused by a lack of PSEN endoproteolysis. **(A,B)** Blue Native PAGE/western blot analysis of γ-secretase complexes. 12 ug of DMM dissolved membranes from fibroblasts containing WT (+/+) or knockout (−/−) alleles of *Psenen* or *Psen1&2* as indicated. *Psenen^−/−^* fibroblasts and *Psen1&Psen2^dKO^* fibroblasts were transduced with the non-cleaved human PSEN1 mutant delta exon 9 (ΔE9). Panel A shows the blot stained with human specific PSEN1 CT antibody (B14) and Panel B shows the same blot reprobed with Mab5232, a mouse and human specific antibody. This shows that PSEN1 ΔE9 is incorporated into the γ-secretase complexes (indicated with full) in the reconstituted *Psen1&2^dKO^* and in the trimeric complex in the absence of PSENEN. * Indicates a non-specific signal at the upper end of the gel. **(C)** *Psenen^−/−^* and *Psen1&2^dKO^* fibroblasts were transduced with PSEN1 ΔE9 as indicated in the figure on the X-axis. The γ-secretase substrates APP containing the Swedish mutation or NotchDE were transduced into the fibroblasts and Aβ_40_, Aβ_42_ and APP CTF or NICD production were assessed by ELISA (Aβ_40,_ blue open circle, and Aβ_42,_ black open square) or by western blot (APPCTF, black circle and NICD,blue square) as indicated in the legend of the graph. None of these γ-secretase products are generated in the *Psen*1&2^dKO^ cells or the *Psenen^−/−^* cells and an accumulation of APP CTFs is observed. Processing of APP and Notch is partially rescued upon PSEN1 ΔE9 expression in *Psen1&2^dKO^* but not in *Psenen^−/−^.* **(D)** AICD (the intracellular domain of APP generated by γ-secretase) is generated in microsomal fractions from wildtype fibroblasts (WT) in the presences of APP C99 substrate and is blocked by the addition of 10µM γ-secretase inhibitor L-685,458 (+GSI). The signal present in Lane 1 indicates the background level of the assay. AICD was absent in cell free reactions using *Psenen^−/−^* microsomal fractions. Expression of human PSEN1 ΔE9 in *Psen1&2^dKO^* restored AICD production, while this is not observed in the *Psenen^−/−^* assay. **(E)** ELISA of Aβ_40_ (black circle) and Aβ_42_ (blue square) of conditioned medium of WT fibroblasts (WT) shows detectable levels of Aβ_40_ signal at 1% of the input material. Thus, Aβ levels in *Psenen^−/−^* fibroblasts are lower than 1% of the Aβ levels in WT fibroblasts. Aβ levels were not detectable in *Psenen^−/−^* fibroblasts rescued with the PS1ΔE9 mutant. Low amounts of Aβ levels in *Ncstn^−/−^* fibroblasts could be detected, in line with previous findings (40), and confirming the sensitivity of the assay.

Since PSENEN is considered to induce PSEN endo-proteolysis, we wondered whether the lack of activity of the trimeric complex in *Psenen^−/−^* fibroblasts was simply the consequence of the fact that PSEN fails to undergo endo-proteolysis in the absence of PSENEN. To investigate this, we stably expressed the PSEN1 delta exon9 (PSEN1 ΔE9) mutant in the *Psenen^−/−^* fibroblasts. PSEN1 ΔE9 is active without the need for endo-proteolytical processing (29, 46, 47). Even though PSEN1 ΔE9 was incorporated in the trimeric complex (Fig. 3A, B), we failed to detect any γ-secretase activity (Fig. 3C and D, lane indicated with *Psenen^−/−^*+ΔE9). In contrast, the PSEN1 ΔE9 mutant rescued γ-secretase activity in fibroblasts lacking PSEN1 and PSEN2, proving that it is indeed active in its non-cleaved form (lane labeled Psen1&2^dKO^ + ΔE9.) This result indicates that PSENEN is not only essential for the endoproteolysis of PSEN but also plays a role in the active γ-secretase complex. Indeed a role of PSENEN in (i) the stabilization of the PSEN fragments after heterodimerization (25), (ii) the specificity of γ-secretase activity (38) and (iii) the accessibility or the affinity of γ-secretase to the substrate (48) has been reported.

Ahn *et al.*, demonstrated that PSEN1 ΔE9 exhibits activity on its own, without the need for other γ-secretase components (34). However, in our experimental conditions, PSEN1 ΔE9 is present in the trimeric complex, which may change the properties of PSEN1 ΔE9. Therefore, we conclude that in the context of the γ-secretase complex, PSENEN is needed for γ-secretase activity beyond endo-proteolysis, considering that our assay conditions would have detected 1% of the wildtype activity.

### SCAM analysis of PSENEN

We decided to use the scanning cysteine accessibility method (SCAM) which has proven to be a valuable approach in the analysis of the structure-function of the PSEN1 subunit of the complex (22–24). We first generated a cysteine-free form of PSENEN (CL PSENEN) by replacing the unique cysteine (C15) in wild type PSENEN by an alanine. The CL PSENEN variant was able to rescue γ-secretase complex formation in the *Psenen^−/−^* fibroblasts as seen in Blue Native PAGE (Fig. 2C, D). PSEN1 cleavage and NCSTN maturation were confirmed using SDS-PAGE/Western blotting (Fig. 2D). Importantly, as judged by APP CTF protein levels, γ-secretase proteolytic activity was restored to the same extent as in *Psenen^−/−^* fibroblasts rescued with wild type PSENEN (Fig. 2D). Therefore, we concluded that CL PSENEN is a suitable backbone for further analysis with SCAM.

We incorporated cysteine residues throughout PSENEN by site directed mutagenesis (see: Fig. 5A). The PSENEN mutants were stably expressed in the *Psenen^−/−^* fibroblasts.

### Glycine 22 and Proline 27 are involved in complex formation and/or stabilization

Two cysteine mutants, i.e., G22C and P27C, did not reconstitute fully γ-secretase levels (Fig 4). Interestingly, although the G22C mutant was expressed to normal levels, complex formation, maturation, and activity were severely compromised (Fig 4). The P27C mutant showed low expression levels (Fig 4B, C) but sufficient to partially restore NCTSN maturation and enzymatic activity (AICD generation in panel 4D). This suggests that the P27C mutant is unstable, but once incorporated into the complex, can replace wild type PSENEN.

**Fig. 4:**
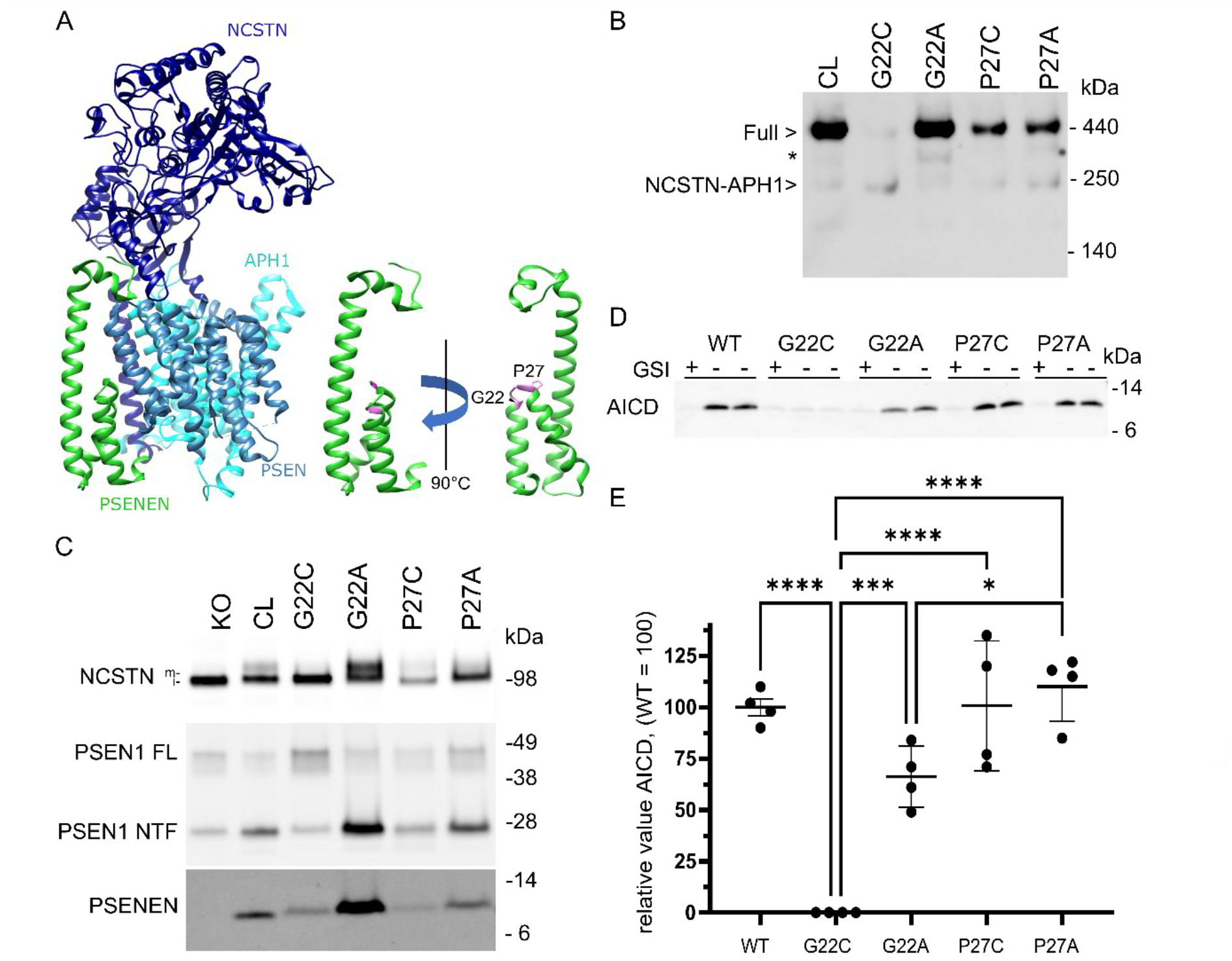
The amino acids G22 and P27 of PSENEN are involved in complex formation and/or stabilization. **(A)** Structural representation of γ-secretase complex and positions of G22 and P27 (pink) in PSENEN (green). **(B)** Blue Native PAGE/western blot analysis of the different *Psenen* mutants. Expression of cysteine less (CL) PSENEN in *Psenen^−/−^* fibroblasts rescues full γ-secretase complex formation (lane 1). In contrast, expression of the G22C mutant compromises complex formation to a large extent (lane 2). Replacement of G22 with alanine (G22A) restores complex assembly (lane 3). Expression of P27C mutant results in a partially rescue of γ-secretase complex (lane 4). Replacement of P27 with an alanine (P27A) has similar effects as the P27C mutation (lane 5). * indicates the detergent induced complex composed of PSEN1 CTF, APH1 and NCSTN (66). **(C)** SDS-PAGE/western blot analysis of *Psenen^−/−^* fibroblasts expressing the G22 and P27 mutants. The PSENEN G22C mutant hardly rescues NCSTN maturation and decreases endo-proteolysis of PSEN1 (lane 3), which agrees with the low levels of full complex seen in Blue Native PAGE analysis (panel A). Replacement of G22 by alanine (lane 4) restores NCSTN maturation and PSEN1 endo-proteolysis to comparable levels as CL PSENEN (lane 2). Mutation of the P27 to cysteine or alanine results in very low PSENEN levels. Nevertheless, the levels of expression are sufficient to restore low levels of γ-secretase complex (lane 5,6). **(D)** Western blot of AICD generated in microsomes isolated from *Psenen^−/−^* fibroblasts transduced with the indicated PSENEN mutants in the presence or absence of 10µM L-685,458 (GSI). Notice that all mutants apart from G22C rescue AICD generation. **(E)** The panel shows quantitation of 4 experiments as shown in D. AICD signal is equal to +GSI condition = background of the assay and AICD levels were normalized to PSEN1 NTF levels to obtain specific γ-secretase activity. Data of 4 experiments are presented in the graph. ****: p< 0,0001, ***p=0.0008, * p=0.0235

High-resolution cryo-EM data for the γ-secretase complex (PDB: 5A63) show that the N-terminus of PSENEN is in the cytosol. Two helical domains (Asn8-Phe23 and Pro27-Leu43) are partially inserted in the membrane forming a U-turn structure and are followed by an intracellular loop (Val44-Gly49), a full membrane-spanning helical domain (Gln50-Tyr81) and a short extracellular C-terminus (Arg82-Pro101). The PSENEN’s C-terminus is located at the interface between NCSTN and PSEN1 in this structure (Fig 4A). Gly22 is located almost in the middle of the membrane at the end of the first partial-transmembrane domain; while Pro27 is present at the U-turn connecting the two partial transmembrane helices (Fig 4A).

Glycine and Prolines are frequent amino acids in transmembrane domains (TMD) of membrane proteins (49), and both contribute to TM dynamics, but their specific roles depend on the local environment (22, 23). Glycine often functions as an interface between individual TM α-helices (50), and when positioned close to a Pro, it may enhance local dynamics and thus affect protein function (51). The structural data show that the conserved Gly22 and Pro27 residues contribute to the formation of the U-turn structure, and thus overall fold of PSENEN. Gly22 enables proximity between the two half-helical structures and the formation of a H-bonding with the backbone of Leu26. To investigate this further we introduced alanine at these two positions. In contrast to G22C, the G22A mutation displayed γ-secretase complex levels comparable to CL PSENEN (Fig. 4C). Furthermore, NCSTN maturation was rescued to a large extent (Fig. 4C) and cell free activity assays demonstrated AICD production (Fig. 4D, E). These results show that the backbone flexibility provided by G22 is not entirely essential for the function of the γ-secretase complex; however, the functional data also indicate that while the short side chain of alanine (G22A) is tolerated, the larger cysteine side chain sterically disturbs helix-helix interactions in the PSENEN fold and thus affects the assembly and/or stability of the protease complex. The P27A mutation, in contrast, had the same effect on PSENEN levels and γ-secretase maturation and activity as the P27C mutation (Fig. 4B). The low levels of mutant PSENEN P27A that assemble into the enzyme complex display similar specific activity as CL PSENEN, indicating that the mutant assembles into functional γ-secretase complexes (Fig. 4D and 4E). Prolines are usually found in irregular structures such as β-turns and α-helical capping motifs (52). In PSENEN, mutation of P27 to another amino acid results in loss of stability of PSENEN and the γ-secretase complex.

### Hydrophobic domain 1 and 2 are water accessible from the extracellular side

To delineate the solvent accessibility of the PSENEN amino acid residues, we tested the reactivity of the introduced cysteines to EZ-linked Biotin-HPDP (HPDP-biotin) (Fig. 5A and 5B). HPDP-biotin is a membrane-permeable sulfhydryl-reactive reagent that can only react with free sulfhydryl groups exposed to a hydrophilic environment. The amino acids Cys15 (WT) and Y18C-G21C, F25C and F28C-W30C, W36C, all present in the partial TM hydrophobic domains 1-2, were water accessible. In contrast, cysteines at positions N33, I34 (see also MTSEA experiment), F37 and R39 did not react with the probe. The results support a model in which part of the first two hairpin forming hydrophobic domains of PSENEN (helix 1 and helix 2) are exposed to a hydrophilic environment. Furthermore, in agreement with the structural data, residues F42C, E49C, Q50C, K54C, W58C connecting the hydrophobic domain 2 and TMD 3, as well as the extracellular W85C were also labelled. In contrast, the A61C to Q79C residues in PSENEN were not labeled by the HPDP-biotin, indicating that this part of the full transmembrane domain (helix 3) is hydrophobic (Fig 5A and 5B).

**Fig. 5:**
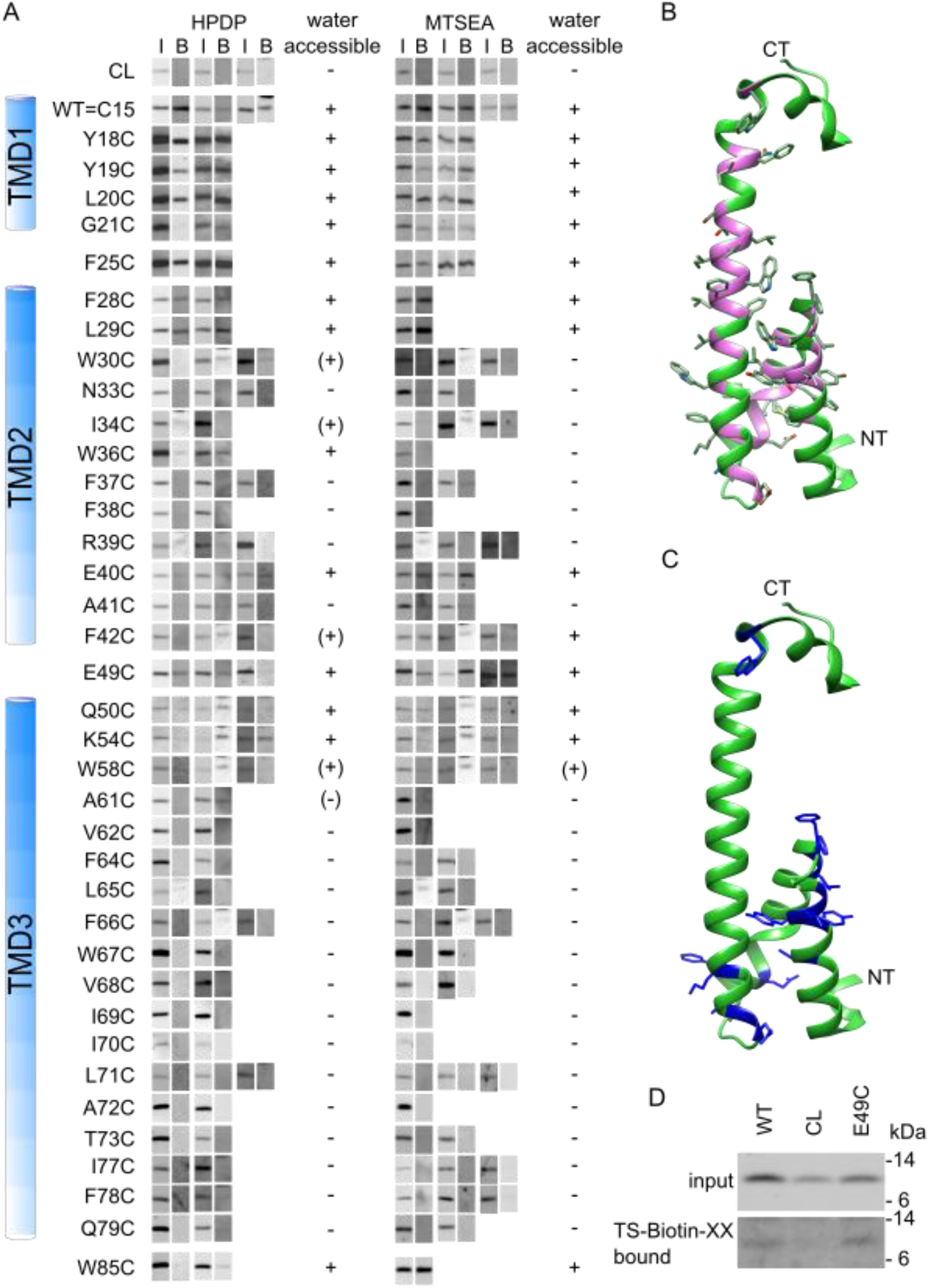
Water accessibility pattern of unique cysteines in PSENEN. **(A)** Intact cells were exposed to the membrane permeable sulfhydryl-specific reagent EZ-linked Biotin-HPDP or the membrane impermeable sulfhydryl-specific reagent MTSEA-biotin as indicated. Biotinylated proteins were precipitated by neutravidin beads in the presence of Triton X-100. Input (I) and bound (B) fractions were separated by SDS-PAGE (4-12% BisTris) and the presence of biotinylated PSENEN was detected using the B126 antibody. Wild type PSENEN (WT) with its luminal cysteine at the N-terminus, was used as a positive control, and cysteine-less PSENEN (CL) was used as a negative control. + and −: indicate reactive and not reactive to the reagent, respectively. The secondary structure of PSENEN is indicated at the left. **(B)** Three-dimensional model of PSENEN. The side chains of the amino acids targeted in the cysteine scan are shown. The purple areas indicate the positions of the amino acids that were targeted in the cysteine scan. **(C)** Three-dimensional model of PSENEN. The positions of amino acids in the backbone of the structure that are exposed to the hydrophilic environment as detected with MTSEA are indicated in blue. **(D)** As a control for the reliability of the MTSEA-reagent, intact cells were treated with the membrane-impermeable TS-Biotin-XX in the same way as in (A). The PSENEN cysteine mutant E49C shows reactivity to the impermeable TS-XX-Biotin, confirming the results of the MTSEA-biotin reagent.

Next, we evaluated the reactivity of cysteines to the membrane-impermeable MTSEA-biotin (Fig. 5A, B). Remarkably, most residues in the partial TDM 1-2 of PSENEN that were labeled by the HPDP-biotin probe were also accessible by the MTSEA-biotin, <u>with the exception of</u> W30C and W36C. Residues F42C, E49C, Q50C and K54C between HD2 and TM3 are labelled by the MTSEA-biotin reagent. The labeling pattern confirms that part of the HD1 and 2, together with the connecting loop and the beginning of TM3 are accessible from the extracellular/luminal site (Fig. 5C). The labelling of Cys49 was confirmed using another membrane-impermeable reagent (TS-XX-biotin) (Fig. 5D), which contains, in contrast to the neutral methanethiosulfonate group of MTSEA-biotin, a negatively charged thiosulfate group and is therefore even less likely to cross the cell membrane. Based on these findings, we speculate that the hydrophobic domains 1 and 2 and the linker with the transmembrane domain 3 are connected to the aqueous environment of the active site in PSEN. The transmembrane domain 3, in contrast, remains largely unlabeled (14 residues tested between A61 and Q79) by both biotin moieties, indicating that this part of PSENEN is embedded in a hydrophobic environment (Fig 5C).

### Crosslinking studies locate E49 and PSEN1 CTF in close proximity

The catalytic site of γ-secretase is situated at the interface between PSEN NTF and PSEN CTF (21–23, 53) in a water-containing cavity. The accessibility results indicate that the N-terminus containing the two hydrophobic domains of PSENEN relates to the catalytic pore in γ-secretase. To investigate this further, we performed bifunctional cross-linking assays involving a cysteine at E49 which is in the U turn linker between the two hydrophobic domains.

Microsomal membrane fractions from the E49C PSENEN mutant were treated with SPDP, a heterobifunctional cross linker with spacer arm of 6.8Å, at 4°C. SPDP conjugates primary amine (mainly lysines) and sulfhydryl (cysteines) groups of proteins. After cross linking, conjugated products were separated by SDS-PAGE in non-reducing conditions and analyzed by western blotting. Antibodies against PSENEN showed a higher mobility band for the PSENEN E49C mutant (Fig. 6, lane 6, arrow). This band was also detected with an antibody against PSEN CTF (Fig. 6, lane 10 arrow), while antibodies against other γ-secretase components (NCSTN, APH1 and PSEN1 NTF) did not react with this band. Moreover, the band was not observed when the free sulfhydryls were blocked with the alkylating agent N-ethylmaleimide (NEM) prior to the cross-linking reaction (Fig. 6 lane 5) or when reducing conditions were applied. No cross-linked products were detected with the cysteine-less (CL) PSENEN (Fig. 6 lane 1-3 and lane 7,8). The molecular weight of the band (~26 kDa) is in accordance with a cross linked product between PSENEN en PSEN1 CTF. Therefore, our results show that the loop of PSENEN and the PSEN1 CTF are at a distance equal or less than 6.8Å from each other in the γ-secretase complex. Considering the available structural data, the crosslink may involve Cys49 in PSENEN and an amine group within the large intracellular (N-terminal) part of PSEN-CTF. This flexible, unstructured region in PSEN-CTF, not seen in the structure, is the only region that could approach the intracellular loop in PSENEN at the short distance reported by the cross-linking experiments. Alternatively, one would need to allude to large conformational changes in PSEN-CTF to bring the loop in PSENEN to this short distance.

**Fig. 6:**
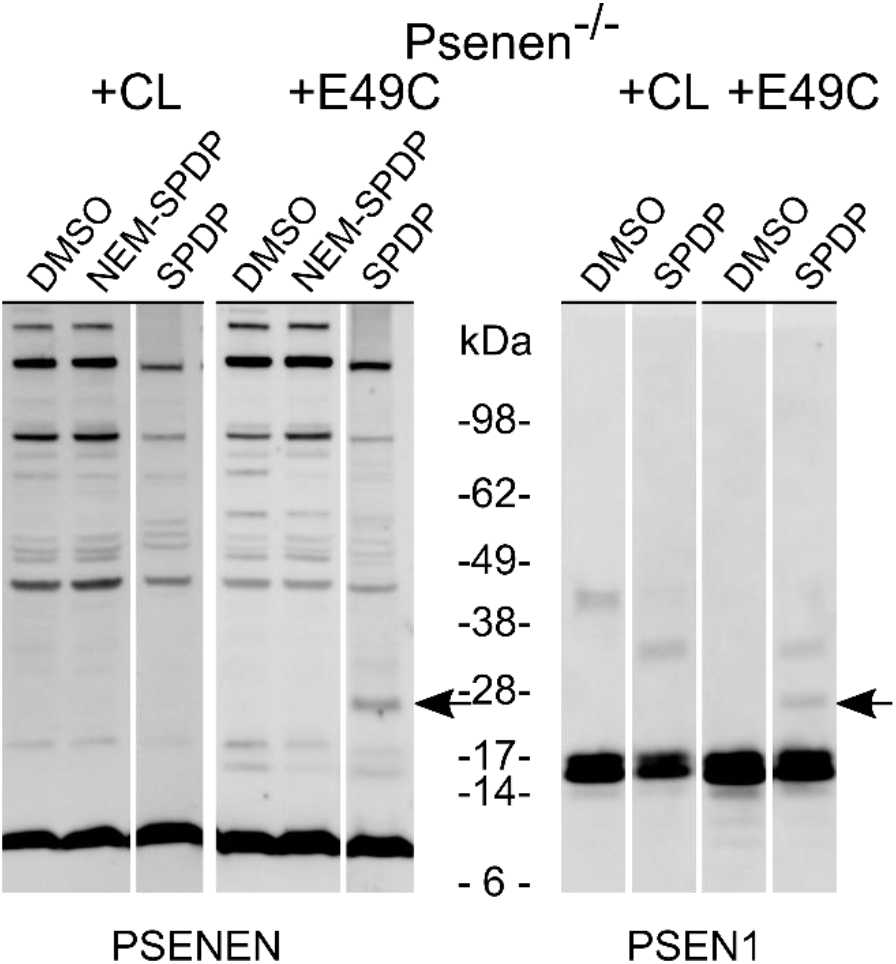
Crosslinking studies reveal close proximity of E49 in PSENEN to PSEN1 CTF. Membrane fractions of *Psenen-^−/−^* fibroblasts transduced with cysteine-less (CL) PSENEN or PSENEN E49C were treated with the heterobifunctional cysteine-amine crosslinker SPDP (spacer arm: 6,8Å). Blocking of the free sulfhydryl groups with NEM was performed as negative control. As additional negative control, crosslinker was omitted in the sample (DMSO). Protein extracts were separated in SDS-PAGE under non-reducing conditions on a 4-12% BisTris gel and visualized in western blot with antibodies against γ-secretase components PSENEN or PSEN1. A band with a molecular weight of approximately 26 kDa was observed with PSENEN and PSEN1 CTF antibody, only in the samples transduced with the E49C mutant in the presence of crosslinker (black arrow).

Until now, only PSENEN and PSEN modifications have been implicated in changes in γ-secretase activity (ratio changes) (15, 38). This implies that PSENEN may play a functional role in the γ-secretase activity, which is clearly corroborated by our experiments shown in figure 3E. While a trimeric γ-secretase complex containing PSENEN but not NCSTN remains partially active (3-6% of the wildtype complex), the trimeric complex containing NCSTN but not PSENEN is completely inactive. Even when we expressed a functional mutant of PSEN1, PSEN1 ΔE9, which does not need endo-proteolytical activation, the trimeric APH1-NCT-PSEN1 ΔE9 complex remained inactive, confirming that PSENEN is needed beyond the endo-proteolysis that activates the enzyme.

Our previous report (18) has been retracted because of doubts raised by the editors regarding some of the figures. We provide here reassembled figures which should restore trust in the conclusions of our previous work. The Notch deficient phenotype of the *Psenen^−/−^* mice ((17, 54–58), the *Psenen^−/−^* fibroblasts to study structure-function (59–62), the role of PSENEN in complex formation (55, 57, 63–65) and the data of the cysteine scan (19, 28, 62) were used and cited in other publications and based on the reported data here, these citations remain fully valid. One major change in the current manuscript is the updated interpretation of the cysteine scan data presented in figure 5A. This interpretation takes fully into account the information regarding structure of the γ-secretase obtained by recent cryo-EM approaches which show that PSENEN makes a reentrant loop and that the N- and C-terminus of PSENEN face opposite sides of the cell membrane (19, 20). This data was not available when we published our original manuscript but were used in the elegant work of Zhao et al (28) which provided a corrected interpretation of the structure. Our data reveal that the hydrophobic domains, helix 1 & 2 in PSENEN are inserted in the membrane but are surrounded by a polar environment. In contrast, the full membrane spanning TMD3 is mostly hydrophobic.

## ACKNOWLEDGEMENTS

We want to thank Galapagos for providing the adenoviral vectors. This work was supported by VIB, FWO, SA0-FRMA, the Federal Office for Scientific Affairs, Belgium (IUAP P6/43), a Methusalem grant of the KU Leuven and the Flemish Government and Memosad (FZ-2007-200611) of the European Union. BDS is supported by the Arthur Bax and Anna Vanluffelen chair for Alzheimer’s disease.

## Author contribution

Conceptualization: BDS, LCG, AT, LB

Methodology Development or design of methodology: all authors

Formal analysis: all authors

Investigation: all authors

Resources: BDS

Data Curation: LS

Writing - Original Draft: LB, BDS

Writing - Review & Editing Preparation: all authors

Visualization: LS

Supervision: BDS, LCG, AT

Funding acquisition: BDS

## Data Availability

The cell lines and raw data described in this paper are available upon request.

## Supporting information

This article does not contain supporting information.

## REFERENCES

1. De Strooper, B. (2003) Aph-1, Pen-2, and Nicastrin with Presenilin generate an active gamma-secretase complex. Neuron. 38, 9–12

2. De Strooper, B., Saftig, P., Craessaerts, K., Vanderstichele, H., Guhde, G., Annaert, W., et al. (1998) Deficiency of presenilin-1 inhibits the normal cleavage of amyloid precursor protein. Nature. 391, 387–390

3. De Strooper, B., Annaert, W., Cupers, P., Saftig, P., Craessaerts, K., Mumm, et al. (1999) A presenilin-1-dependent g-secretase-like protease mediates release of Notch intracellular domain. Nature 398, 518–522

4. Struhl, G., and Greenwald, I. (1999) Presenilin is required for activity and nuclear access of Notch in Drosophila. Nature. 398, 522–5

5. Levitan, D., and Greenwald, I. (1995) Facilitation of lin-12-mediated signaling by sel-12, a Caenorhabditis elegans S182 Alzheimer’s disease gene. Nature. 377, 351–354

6. Goutte, C., Hepler, W., Mickey, K. M., and Priess, J. R. (2000) aph-2 encodes a novel extracellular protein required for GLP-1-mediated signaling. Development. 127, 2481– 92

7. Herreman, A., Hartmann, D., Annaert, W., Saftig, P., Craessaerts, K., Serneels, L., et al. (1999) Presenilin 2 deficiency causes a mild pulmonary phenotype and no changes in amyloid precursor protein processing but enhances the embryonic lethal phenotype of presenilin 1 deficiency. Proc Natl Acad Sci U S A. 96, 11872–7

8. Serneels, L., Dejaegere, T., Craessaerts, K., Horré, K., Jorissen, E., Tousseyn, T., et al. (2005) Differential contribution of the three Aph1 genes to gamma-secretase activity in vivo. Proc Natl Acad Sci U S A. 102, 1719–24

9. Li, J., Fici, G. J., Mao, C.-A., Myers, R. L., Shuang, R., Donoho, G. P., et al. (2003) Positive and negative regulation of the gamma-secretase activity by nicastrin in a murine model. J Biol Chem. 278, 33445–9

10. Donoviel, D. B., Hadjantonakis, A. K., Ikeda, M., Zheng, H., Hyslop, P. S., and Bernstein, A. (1999) Mice lacking both presenilin genes exhibit early embryonic patterning defects. Genes Dev. 13, 2801–10

11. Francis, R., McGrath, G., Zhang, J., Ruddy, D. A., Sym, M., Apfeld, J., et al. (2002) aph-1 and pen-2 are required for Notch pathway signaling, gamma-secretase cleavage of betaAPP, and presenilin protein accumulation. Dev Cell. 3, 85–97

12. Huppert, S. S., Le, A., Schroeter, E. H., Mumm, J. S., Saxena, M. T., Milner, L. A., et al. (2000) Embryonic lethality in mice homozygous for a processing-deficient allele of Notch1. Nature. 405, 966–70

13. Wong, P. C., Zheng, H., Chen, H., Becher, M. W., Sirinathsinghji, D. J., Trumbauer, M. E., et al. (1997) Presenilin 1 is required for Notch1 and DII1 expression in the paraxial mesoderm. Nature. 387, 288–92

14. Shen, J., Bronson, R. T., Chen, D. F., Xia, W., Selkoe, D. J., and Tonegawa, S. (1997) Skeletal and CNS defects in Presenilin-1-deficient mice. Cell. 89, 629–639

15. Uemura, K., Lill, C. M., Li, X., Peters, J. A., Ivanov, A., Fan, Z., et al. (2009) Allosteric modulation of PS1/gamma-secretase conformation correlates with amyloid beta(42/40) ratio. PLoS One. 4, e7893

16. Ma, G., Li, T., Price, D. L., and Wong, P. C. (2005) APH-1a is the principal mammalian APH-1 isoform present in gamma-secretase complexes during embryonic development. J Neurosci. 25, 192–8

17. Jurisch-Yaksi, N., Sannerud, R., and Annaert, W. (2013) A fast growing spectrum of biological functions of γ-secretase in development and disease. Biochim Biophys Acta. 1828, 2815–27

18. Bammens, L., Chávez-Gutiérrez, L., Tolia, A., Zwijsen, A., and De Strooper, B. (2011) Functional and topological analysis of Pen-2, the fourth subunit of the gamma-secretase complex. J Biol Chem. 286, 12271–82

19. Lu, P., Bai, X. C., Ma, D., Xie, T., Yan, C., Sun, L., Y., et al. (2014) Three-dimensional structure of human γ-secretase. Nature. 512, 166–170

20. Bai, X. C., Yan, C., Yang, G., Lu, P., Ma, D., Sun, L., et al. (2015) An atomic structure of human γ-secretase. Nature. 525, 212–217

21. Wolfe, M. S., Xia, W., Ostaszewski, B. L., Diehl, T. S., Kimberly, W. T., and Selkoe, D. J. (1999) Two transmembrane aspartates in presenilin-1 required for presenilin endoproteolysis and gamma-secretase activity. Nature. 398, 513–7

22. Sato, C., Morohashi, Y., Tomita, T., and Iwatsubo, T. (2006) Structure of the catalytic pore of gamma-secretase probed by the accessibility of substituted cysteines. J Neurosci. 26, 12081–8

23. Tolia, A., Chávez-Gutiérrez, L., and De Strooper, B. (2006) Contribution of presenilin transmembrane domains 6 and 7 to a water-containing cavity in the gamma-secretase complex. J Biol Chem. 281, 27633–42

24. Tolia, A., and De Strooper, B. (2009) Structure and function of gamma-secretase. Semin Cell Dev Biol. 20, 211–8

25. Petit, D., Hitzenberger, M., Lismont, S., Zoltowska, K. M., Ryan, N. S., Mercken, M., et al. (2019) Extracellular interface between APP and Nicastrin regulates Aβ length and response to γ-secretase modulators. EMBO J. 10.15252/embj.2019101494

26. Crystal, A. S., Morais, V. A., Pierson, T. C., Pijak, D. S., Carlin, D., Lee, V. M.-Y., et al. (2003) Membrane topology of gamma-secretase component PEN-2. J Biol Chem. 278, 20117–23

27. Bergman, A., Hansson, E. M., Pursglove, S. E., Farmery, M. R., Lannfelt, L., Lendahl, U., et al. (2004) Pen-2 is sequestered in the endoplasmic reticulum and subjected to ubiquitylation and proteasome-mediated degradation in the absence of presenilin. J Biol Chem. 279, 16744–53

28. Zhang, X., Yu, C. J., and Sisodia, S. S. (2015) The topology of pen-2, a γ-secretase subunit, revisited: Evidence for a reentrant loop and a single pass transmembrane domain. Mol Neurodegener. 10.1186/s13024-015-0037-4

29. Thinakaran, G., Borchelt, D. R., Lee, M. K., Slunt, H. H., Spitzer, L., Kim, G., et al. (1996) Endoproteolysis of presenilin 1 and accumulation of processed derivatives in vivo. Neuron. 17, 181–90

30. Luo, W., Wang, H., Li, H., Kim, B. S., Shah, S., Lee, H.-J., T, et al. (2003) PEN-2 and APH-1 coordinately regulate proteolytic processing of presenilin 1. J Biol Chem. 278, 7850–4

31. Kim, S.-H., and Sisodia, S. S. (2005) A sequence within the first transmembrane domain of PEN-2 is critical for PEN-2-mediated endoproteolysis of presenilin 1. J Biol Chem. 280, 1992–2001

32. Takasugi, N., Tomita, T., Hayashi, I., Tsuruoka, M., Niimura, M., Takahashi, Y., et al. (2003) The role of presenilin cofactors in the gamma-secretase complex. Nature. 422, 438–41

33. Watanabe, N., Tomita, T., Sato, C., Kitamura, T., Morohashi, Y., and Iwatsubo, T. (2005) Pen-2 is incorporated into the gamma-secretase complex through binding to transmembrane domain 4 of presenilin 1. J Biol Chem. 280, 41967–75

34. Ahn, K., Shelton, C. C., Tian, Y., Zhang, X., Gilchrist, M. L., Sisodia, S. S., et al. (2010) Activation and intrinsic gamma-secretase activity of presenilin 1. Proc Natl Acad Sci U S A. 107, 21435–40

35. Prokop, S., Haass, C., and Steiner, H. (2005) Length and overall sequence of the PEN-2 C-terminal domain determines its function in the stabilization of presenilin fragments. J Neurochem. 94, 57–62

36. Hasegawa, H., Sanjo, N., Chen, F., Gu, Y.-J., Shier, C., Petit, A., et al. (2004) Both the sequence and length of the C terminus of PEN-2 are critical for intermolecular interactions and function of presenilin complexes. J Biol Chem. 279, 46455–63

37. Prokop, S., Shirotani, K., Edbauer, D., Haass, C., and Steiner, H. (2004) Requirement of PEN-2 for stabilization of the presenilin N-/C-terminal fragment heterodimer within the gamma-secretase complex. J Biol Chem. 279, 23255–61

38. Isoo, N., Sato, C., Miyashita, H., Shinohara, M., Takasugi, N., Morohashi, Y., et al. (2007) Abeta42 overproduction associated with structural changes in the catalytic pore of gamma-secretase: common effects of Pen-2 N-terminal elongation and fenofibrate. J Biol Chem. 282, 12388–96

39. Kounnas, M. Z., Danks, A. M., Cheng, S., Tyree, C., Ackerman, E., Zhang, X., et al. (2010) Modulation of gamma-secretase reduces beta-amyloid deposition in a transgenic mouse model of Alzheimer’s disease. Neuron. 67, 769–80

40. Zhao, G., Liu, Z., Ilagan, M. X. G., and Kopan, R. (2010) γ-secretase composed of PS1/Pen2/Aph1a can cleave notch and amyloid precursor protein in the absence of nicastrin. Journal of Neuroscience. 30, 1648–1656

41. Akabas, M. H., Stauffer, D. A., Xu, M., and Karlin, A. (1992) Acetylcholine receptor channel structure probed in cysteine-substitution mutants. Science. 258, 307–10

42. Annaert, W. G., Esselens, C., Baert, V., Boeve, C., Snellings, G., Cupers, P., et al. (2001) Interaction with Telencephalin and the Amyloid Precursor Protein Predicts a Ring Structure for Presenilins. Neuron 20. 579–589

43. Esselens, G., Oorschot, V., Baert, V., Raemaekers, T., Spittaels, K., Serneels, et al. (2004) Presenilin 1 mediates the turnover of telencephalin in hippocampal neurons via an autophagic degradative pathway. Journal of Cell Biology. 166, 1041–1054

44. Kakuda, N., Funamoto, S., Yagishita, S., Takami, M., Osawa, S., Dohmae, N., et al. (2006) Equimolar production of amyloid β-protein and amyloid precursor protein intracellular domain from β-carboxyl-terminal fragment by γ-secretase. Journal of Biological Chemistry. 281, 14776–14786

45. Steiner, H., Winkler, E., Edbauer, D., Prokop, S., Basset, G., Yamasaki, A., et al. (2002) PEN-2 is an integral component of the gamma-secretase complex required for coordinated expression of presenilin and nicastrin. J Biol Chem. 277, 39062–5

46. Prihar, G., Verkkoniem, A., Perez-Tur, J., Crook, R., Lincoln, S., Houlden, H., et al. (1999) Alzheimer disease PS-1 exon 9 deletion defined. Nat Med. 5, 1090

47. Perez-Tur, J., Froelich, S., Prihar, G., Crook, R., Baker, M., Duff, K., et al. (1995) A mutation in Alzheimer’s disease destroying a splice acceptor site in the presenilin-1 gene. Neuroreport. 7, 297–301

48. Shiraishi, H., Sai, X., Wang, H.-Q., Maeda, Y., Kurono, Y., Nishimura, M., et al. (2004) PEN-2 enhances gamma-cleavage after presenilin heterodimer formation. J Neurochem. 90, 1402–13

49. Baker, J. A., Wong, W.-C., Eisenhaber, B., Warwicker, J., and Eisenhaber, F. (2017) Charged residues next to transmembrane regions revisited: “Positive-inside rule” is complemented by the “negative inside depletion/outside enrichment rule”. BMC Biol. 15, 66

50. Javadpour, M. M., Eilers, M., Groesbeek, M., and Smith, S. O. (1999) Helix packing in polytopic membrane proteins: role of glycine in transmembrane helix association. Biophys J. 77, 1609–18

51. Jacob, J., Duclohier, H., and Cafiso, D. S. (1999) The role of proline and glycine in determining the backbone flexibility of a channel-forming peptide. Biophys J. 76, 1367–76

52. Parker, M. H., and Hefford, M. A. (1997) A consensus residue analysis of loop and helix-capping residues in four-alpha-helical-bundle proteins. Protein Eng. 10, 487–96

53. Nyabi, O., Bentahir, M., Horré, K., Herreman, A., Gottardi-Littell, N., Van Broeckhoven, C., et al. (2003) Presenilins Mutated at Asp-257 or Asp-385 Restore Pen-2 Expression and Nicastrin Glycosylation but Remain Catalytically Inactive in the Absence of Wild Type Presenilin. Journal of Biological Chemistry. 278, 43430–43436

54. Cheng, S., Liu, T., Hu, Y., Xia, Y., Hou, J., Huang, C., Z, et al.(2019) Conditional Inactivation of Pen-2 in the Developing Neocortex Leads to Rapid Switch of Apical Progenitors to Basal Progenitors. J Neurosci. 39, 2195–2207

55. Klein, M., Kaleem, A., Oetjen, S., Wünkhaus, D., Binkle, L., Schilling, S., et al. (2022) Converging roles of PSENEN/PEN2 and CLN3 in the autophagy-lysosome system. Autophagy. 18, 2068–2085

56. Bi, H.-R., Zhou, C.-H., Zhang, Y.-Z., Cai, X.-D., Ji, M.-H., Yang, J.-J., et al. (2021) Neuron-specific deletion of presenilin enhancer2 causes progressive astrogliosis and age-related neurodegeneration in the cortex independent of the Notch signaling. CNS Neurosci Ther. 27, 174–185

57. Xia, Y., Zhang, Y., Xu, M., Zou, X., Gao, J., Ji, M.-H., et al. (2022) Presenilin enhancer 2 is crucial for the transition of apical progenitors into neurons but into not basal progenitors in the developing hippocampus. Development. 10.1242/dev.200272

58. Watanabe, H., Imaizumi, K., Cai, T., Zhou, Z., Tomita, T., and Okano, H. (2021) Flexible and Accurate Substrate Processing with Distinct Presenilin/γ-Secretases in Human Cortical Neurons. eNeuro. 10.1523/ENEURO.0500-20.2021

59. Wouters, R., Michiels, C., Sannerud, R., Kleizen, B., Dillen, K., Vermeire, W., et al. (2021) Assembly of γ-secretase occurs through stable dimers after exit from the endoplasmic reticulum. J Cell Biol. 10.1083/jcb.201911104

60. Holmes, O., Paturi, S., Wolfe, M. S., and Selkoe, D. J. (2014) Functional analysis and purification of a Pen-2 fusion protein for γ-secretase structural studies. J Neurochem. 131, 94–100

61. Hu, C., Zeng, L., Li, T., Meyer, M. A., Cui, M.-Z., and Xu, X. (2016) Nicastrin is required for amyloid precursor protein (APP) but not Notch processing, while anterior pharynx-defective 1 is dispensable for processing of both APP and Notch. J Neurochem. 136, 1246–1258

62. Holmes, O., Paturi, S., Selkoe, D. J., and Wolfe, M. S. (2014) Pen-2 is essential for γ-secretase complex stability and trafficking but partially dispensable for endoproteolysis. Biochemistry. 53, 4393–406

63. Bursavich, M. G., Harrison, B. A., and Blain, J. F. (2016) Gamma secretase modulators: New Alzheimer’s drugs on the horizon? J Med Chem. 59, 7389–7409

64. Angulo-Rojo, C., Manning-Cela, R., Aguirre, A., Ortega, A., and López-Bayghen, E. (2013) Involvement of the Notch pathway in terminal astrocytic differentiation: role of PKA. ASN Neuro. 5, e00130

65. Gertsik, N., Chiu, D., and Li, Y.-M. (2014) Complex regulation of γ-secretase: from obligatory to modulatory subunits. Front Aging Neurosci. 6, 342

66. Fraering, P. C., LaVoie, M. J., Ye, W., Ostaszewski, B. L., Kimberly, W. T., Selkoe, D. J., et al. (2004) Detergent-dependent dissociation of active gamma-secretase reveals an interaction between Pen-2 and PS1-NTF and offers a model for subunit organization within the complex. Biochemistry. 43, 323–33

